# Analysis of genetic control and QTL mapping of essential wheat grain quality traits in a recombinant inbred population

**DOI:** 10.1101/361329

**Authors:** Sonia Goel, Kalpana Singh, Balwant Singh, Sapna Grewal, Neeta Dwivedi, Abdulaziz A. Alqarawi, Elsayed Fathi Abd_Allah, N. K. Singh, Parvaiz Ahmad

## Abstract

Wheat cultivars are genetically crossed for improving end use quality for apt traits as per need of baking industry and broad consumer’s preferences. The processing and baking qualities of bread wheat underlie into genetic make-up of a variety and influence by environmental factors and their interactions. WL711 and C306 derived recombinant inbred lines (RILs) population of 206 was used for phenotyping of quality related traits in three different environmental conditions. The genetic analysis of quality traits showed considerable variation for measurable quality traits with normal distribution and transgressive segregation across the years. From the 206 RIL, few RILs found to be superior to those of the parental cultivars for key quality traitsindicating their potential usefor improvement of end use quality and also suggestingprobability of finding new alleles and allelic combinations from the RIL population. A genetic linkage map including 346 markers was constructed withtotal map distance of 4526.8cM andinterval distance between adjacent markersof 12.9cM. Mapping analysis identified 38 putative QTLs for 13 quality related traits with QTLs explaining 7.9% - 16.8% phenotypic variation spanning over 14 chromosomes i.e. 1A, 1B, 1D, 2A, 2D, 3B, 3D, 4A, 4B, 4D, 5D, 6A, 7A and 7B. Major novel QTLs regions for quality traits have been identified on several chromosome in studied RIL population posing their potential role in marker assisted selection for better bread making quality after validation.

## Introduction

Recent research in wheat contributes to yield enhancement and disease resistance but quality is lacking behind today’s scenario. However, wide consumer demandforced wheat breeders to focus on wheat quality improvement as per consumer’s preference and industrial need. Bread wheat (*Triticum aestivum*L.) is the globally accepted food crop and consumed mainly as baked products. The end-use quality of wheat is governed by plethora of genes network that are majorly affected by environmental conditions. Further, it is measured by its rheological traits like gluten strength, sodium dodecyl sulphate-sedimentation test (SDS sedimentation), farinograph. Quantitative traits loci (QTLs)for quality traits including grain protein content(1-3), grain hardness (4, 5), and dough quality traits namely mixing tolerance, mixing time, dough extensibility and dough tenacity (6, 7) have been mapped. Groos *et al.* (1) reportedfour QTLs for grain protein content (GPC) on chromosomes 2A, 3A, 4D, and 7D.

Linkage mapping and subsequent QTLs mapping is the prerequisite for applying successful marker assisted selection (MAS)program for individual trait. Earlier, MAS was executed in hexaploid wheat for high grain protein content (*Gpc*-B1) which was mapped and introgressed from the wild tetraploid wheat *T. turgidum var. Dicoccoides* (8). Further, role of the QTLs (*Gpc*-B1) for increased GPC was approved in tetraploid and hexaploid wheat using near-isogenic lines (NILs) with distinct *Gpc*-*B1* alleles (9). In additions, two independent studies conducted by Kumar *et al.* (10) and Tabbita *et al.* (11)showed that GPC was increased in Indian and Argentine hexaploid wheat carrying *Gpc*-*B1*. However, pleotropic effect of the QTL *Gpc*-*B1* is associated with reduced grain size and grain yield that ultimately leads to reduction of wheat production (11, 12).

Dough rheological properties and kernel hardness strongly effect end-use quality of wheat. Dough making properties are often used as indicator of thefood baking quality. Dough strength and starch pasting characteristics are reported as quantitative traits therefore their expression isgoverned by multiple genes (13). In the present scenario, no specific bread making quality trait controlling genes have been identified that have direct association with end product quality. Nonetheless,few QTLs for the end productquality traits have been reported (14). Wheat quality was affected by temperature and humidity but their effect isspecific to developmental growth stage. Nuttall *et al.*(15)have been reported that high temperatures during grain filling was responsible for reduced dough strength. Further, Cavanagh *et al*. (16)identified additional traits such as percentage unextractable polymeric protein (% UPP) and dough strength whichwas directly affected by temperature during grain filling stage. The kernel hardness (KH) of wheat grain is a major determinant of endfood productquality. KH refers to the texture of the grain (caryopsis) that represents physical hardness or softness of the endosperm. Kernel hardness is predominantly controlled by the Puroindoline (*Pin*) genes, *Pin a* and *Pin b*, which are part of only the D sub-genome and located on chromosome 5 at the Hardness (*Ha*) locus. Furthermore, different classes’ grain texture has been determined by unique allelic blends of Pin genes (Pin a and Pinb) in wheat with diverse end-use characteristics (17). The key role of the *Pin a* and *Pin b* genes is to determine the structure of the proteins in wheat grain and also possible antimicrobial effects (18). Therefore, to develop a variety with desired kernel hardness pronounced understanding of the allelic composition of *Pin* genes in a diverse set of germplasm is utmost important for the donor parental selection.

In the present study, total 206 RIL population was used for phenotyping of quality related traits in three different locations in India namely Delhi, Karnal and Indore. The aim of the present research was to unravel the genetic factors controlling bread making quality related traits by using wheat mapping population grown in three different environmental conditions through mapping of QTLs associated with these quality traits.

## Materials and methods

### Plant materials and experiment design

In the present study, a mapping population of 206 RILs (F_14:15_) was genotyped and evaluated for different quality traits. The RIL population was developed by crossing of two wheat cultivars ‘WL711’ which was itself developed from S308/Chris/Kalyansona and ‘C306’ developed from RGB/CSL3//2/C591/3/C217/N14//C281 (19). The grain samples were multiplied in the bulk in three independent field experiments conducted at Directorate of Wheat Research (DWR) in 2008, Karnal (76°09’E, 29°60’N; 228.6 M.S.L) (KL08), National Research Centre on Soybean (NRCS) Indore (75°50’E, 22°43’N; 529.9 M.S.L) in 2009 (IN09), and Division of Genetics, Indian Agriculture Research Institute, New Delhi (77°12’E, 28°40’N; 228.6 M.S.L), India in 2010 (DL10). These three regions are geographically located in the tradition wheat agroecosystems. RILs along with parents were sown in three environments in a randomized complete block design (RCBD) pattern with replication in the field. Sowing was done in a plot of 3 rows with 1.5 m long and each row equally spaced by 25 cm and in each row total 30 seeds wereplanted. The RIL was sown in during mid-November and harvesting was done in April at Delhi (DL10) and while in Karnal (KL08) it was sown in early Novemberand harvesting was done in early March at Indore (IN09).

### Quality traits analysis

RILs grain samples collected from each experimental location were analysed at Cereal Quality Laboratory, Division of Genetics, Indian Agriculture Research Institute, New Delhi, India. Samples were hand cleaned and air-aspirated for removing foreign material and shrivelled kernels. The estimation of GPC was done by near-infrared reflectance (NIR) (RACI-CCD, 2010) by using a NIR instrument (Foss 6500, FOSS NIR Systems, Inc., Laurel, MD)(20). Estimated Sedimentation Volume represented in height (mm) of the sediment measured during SDS sedimentation testwas estimated as gluten strength (21). Wheat flour of 206 RILs and the two parental genotypes (WL711 and C306) used for quality analysis was produced by Cyclotec Mill (Tecator AB, Sweden) fitted with a 1 mm sieve. Five flour quality traits namely, breakdown time (BDT), dough development time (DDT), dough stability time (DST), flour water absorption (FWA), and mixing tolerance index (MTI) were recorded by Farinograph (Brabender, Germany) according to AACC 2000 (20).

Clean samples of 20 g seeds having grain moisture content ranged between 10% to 11% was used for analysis of kernel hardness (KH), thousand kernel weight (TKW) and kernel diameter using Single Kernel Characterization system (SKCS) 4100 (Perten Instruments, Australia) using AACC method (2000). Hectolitre weight (HW) was measured as the volume of grain per unit. Further, grain protein gluten was measured as wet and dry gluten using Glutomatic 2200 (Perten Instruments) according to AACC method (2000).

### Statistical analysis of the traits

Statistical and genetic analysis for quality traits was performed byGenStat14 (22). The analysis was conducted in two stageswhile taking account of experimental design factors, firstly spatial analysis (23) to find out best linear unbiased estimates (BLUEs), Broad sense heritability (*h*_B^2^_) value was calculated for each trait across environment as

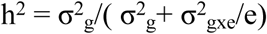

Where σ^2^ _g_= [MS_RIL_ - MS_RILxe_)/e], σ^2^ _gxe_ = MS_RILxe_,

e is the number of environments and MS is the mean square

## Genetic analysis of the traits

Genotyping of the RIL population and linkage map was published in Shukla *et al*. (24).

### Wheat-rice comparison

Candidate genes underlying QTLs of interest were studied by comparing the chromosomal sequences of 1B and 5D chromosomes. The QTLs spanning markers were blast with the genomic DNA sequences of rice (*Oryza sativaL*.) available at TIGR (http://www.tigr.org, http://ncbi.nlm.nih.gov). BLASTN query was performed taking e-value <1 × 10-^19^ and score >100 for comparing the DNA sequences. In order to find out wheat orthologs of the candidate genes their protein sequences werecompared against wheat ESTs (e-value <1 × 10^-25^).

## Results

### Phenotypic data and correlation analysis

Experiments were conducted at three locations in three different years. Performance of both the parents was observed along with RIL populations. WL711 showed low quality score at KL08 comparative to other locations however, C306 showed better performance for traits like SD, FWA, BDT and KH at the same locations and years (Table 1). Measurable phenotypic variation was observed among both the parents for SDS, TKW, WGC, DGC, FWA, DST, MTI, BDT and KH. All quality related traits significantly different extensively among the RILs which were normally distributed and exhibited transgressive segregation (Table 1). GPC and HW showed high broad sense heritability while TKW and KH showed moderate heritability. Highly significant positive correlation were recorded between GPC and WGC,DGC and FWA; between SDS and WGC, DGC, DDT and DST; between DDT and BDT, KH; between DST and BDT. Highly significant but negative correlation were recorded between GPC and TKW; WGC and DST; between MTI and DST, BDT (Table 2).

**Table 1.** Quality parameters in parents and RIL population derived from WL711/C306.

| Traits | Experiment<br>s | Parental lines |  | RIL population |  |  |  |  |
| --- | --- | --- | --- | --- | --- | --- | --- | --- |
|  |  | WL711 | C306 | Min | Max | Mean | CV<br>% | Broad<br>sense<br>heritability |
| Grain protein<br>content (GPC,<br>%) | DL09 | 11.4 ± 0.71 | 14.7 ± 0.16 | 10.5 | 18.5 | 13.9 ± 0.33 | 5.9 | 0.75 |
|  | KL08 | 11.2 ± 0.37 | 13.9 ± 0.24 | 9.8 | 16.2 | 11.5 ± 0.11 | 3.7 | 0.52 |
|  | IN09 | 11.9 ± 0.25 | 15.6 ± 0.21 | 11.2 | 19.9 | 15.9 ± 0.41 | 4.2 | 0.64 |
| Sedimentatio<br>n rate (SDS) | DL09 | 45.8 ± 2.9 | 56.5 ± 1.7 | 24 | 75.8 | 46.0 ± 2.2 | 4.7 | 0.58 |
|  | KL08 | 42.4 ± 1.5 | 53.5 ± 1.1 | 21.6 | 71.5 | 43.0 ± 1.2 | 2.4 | 0.54 |
|  | IN09 | 48.2 ± 1.8 | 57.8 ± 1.3 | 23.7 | 73.5 | 49.0 ± 3.2 | 3.9 | 0.55 |
| Hectolitre<br>weight (HW,<br>g) | DL09 | 77.5 ± 3.6 | 73.8 ± 4.3 | 55.5 | 83.8 | 78.7 ± 2.0 | 6.8 | 0.76 |
|  | KL08 | 72.1 ± 2.4 | 74.1 ± 2.7 | 54.9 | 81.2 | 74.2 ± 1.3 | 4.2 | 0.74 |
|  | IN09 | 79.5 ± 4.1 | 76.5 ± 3.9 | 59.3 | 87.6 | 79.4 ± 4.8 | 3.6 | 0.69 |
| 1000-kernel<br>weight<br>(TKW, g) | DL09 | 35.6 ± 1.4 | 41.5 ± 1.3 | 27.4 | 52.7 | 41.1 ± 1.9 | 3.5 | 0.63 |
|  | KL08 | 33.3 ± 2.1 | 44.2 ± 1.5 | 22.2 | 55.3 | 39.5 ± 1.2 | 3.2 | 0.68 |
|  | IN09 | 37.2 ± 2.7 | 45.5 ± 1.8 | 25.7 | 58.1 | 46.2 ± 1.7 | 3.4 | 0.62 |
| Seed diameter<br>(SD) | DL09 | 2.75 ± 0.18 | 2.98 ± 0.25 | 2.12 | 3.5 | 3.03 ± 1.0 | 9.5 | 0.53 |
|  | KL08 | 2.22 ± 0.14 | 3.18 ± 0.19 | 2.63 | 2.5 | 2.92 ± 1.4 | 8.3 | 0.59 |
|  | IN09 | 2.64 ± 0.19 | 2.76 ± 0.32 | 2.81 | 3.9 | 3.66 ± 1.7 | 9.8 | 0.61 |
| Wet gluten<br>content<br>(WGC, %) | DL09 | 28.5 ± 1.1 | 36.8 ± 1.6 | 23.5 | 48 | 35.5 ± 1.5 | 8.5 | 0.49 |
|  | KL08 | 24.9 ± 1.5 | 37.2 ± 1.3 | 21.2 | 45.9 | 35.8 ± 1.8 | 8.2 | 0.42 |
|  | IN09 | 26.2 ± 1.9 | 39.1 ± 1.4 | 25.7 | 47.3 | 33.3 ± 2.1 | 9.6 | 0.45 |
| Dry gluten<br>content<br>(DGC, %) | DL09 | 9.5 ± 0.6 | 12.5 ± 0.8 | 7 | 14.8 | 11.2 ± 1.2 | 7.4 | 0.41 |
|  | KL08 | 8.9 ± 0.2 | 13.2 ± 0.3 | 5.5 | 13.4 | 10.7 ± 1.3 | 7.2 | 0.39 |
|  | IN09 | 9.8 ± 0.8 | 13.9 ± 0.6 | 8.3 | 14.9 | 11.9 ± 1.7 | 8.1 | 0.46 |
| Flour water<br>absorption<br>(FWA, %) | DL09 | 58.5 ± 1.3 | 64.2 ± 2.4 | 54.6 | 68.5 | 60.8 ± 2.7 | 2.1 | 0.59 |
|  | KL08 | 56.9 ± 1.2 | 66.1 ± 2.2 | 54.8 | 66.2 | 61.1 ± 2.8 | 2.7 | 0.57 |
|  | IN09 | 59.1 ± 1.7 | 66.9 ± 2.8 | 55.2 | 65.9 | 60.9 ± 2.9 | 2.4 | 0.51 |
| Dough<br>development<br>time (DDT,<br>min) | DL09 | 3.8 ± 0.4 | 5.4 ± 0.8 | 2.1 | 8.5 | 4.5 ± 0.6 | 6.4 | 0.55 |
|  | KL08 | 3.3 ± 0.1 | 5.3 ± 0.5 | 2.6 | 8.2 | 4.2 ± 0.2 | 6.2 | 0.49 |
|  | IN09 | 3.5 ± 0.8 | 5.8 ± 0.6 | 2.2 | 8.8 | 4.7 ± 0.8 | 6.7 | 0.61 |
| Dough<br>stability time<br>(DST, min) | DL09 | 2.8 ± 0.2 | 8.5 ± 0.5 | 1.5 | 10.8 | 4.0 ± 0.3 | 6.8 | 0.51 |
|  | KL08 | 2.4 ± 0.3 | 8.3 ± 0.6 | 1.3 | 10.3 | 4.3 ± 0.6 | 6.2 | 0.58 |
|  | IN09 | 2.7 ± 0.8 | 8.2 ± 0.8 | 1.8 | 10.6 | 4.8 ± 0.5 | 6.7 | 0.52 |
| Mixing<br>tolerance<br>index (MSI,<br>F.U) | DL09 | 105.4 ± 6.8 | 45.0 ± 3.2 | 25 | 151.6 | 80.5 ± 2.8 | 5.4 | 0.57 |
|  | KL08 | 102.1 ± 5.2 | 43.0 ± 4.1 | 22.7 | 153.1 | 80.1 ± 2.7 | 5.1 | 0.63 |
|  | IN09 | 107.7 ± 6.5 | 48.0 ± 3.6 | 25.9 | 154.9 | 82.5 ± 2.1 | 5.8 | 0.61 |
| Break down<br>time (BDT,<br>min) | DL09 | 4.2 ± 0.5 | 11.5 ± 0.8 | 2.2 | 14.5 | 7.5 ± 0.6 | 0.8 | 0.34 |
|  | KL08 | 3.1 ± 0.2 | 12.8 ± 0.3 | 2.9 | 12.6 | 7.1 ± 1.2 | 0.2 | 0.38 |
|  | IN09 | 3.8 ± 0.6 | 11.9 ± 0.1 | 2.4 | 15.8 | 7.7 ± 0.8 | 0.9 | 0.35 |
| Kernel<br>hardness<br>(KH) | DL09 | 89 ± 2.5 | 60 ± 3.5 | 99 | 68 | 83.5 ± 4.0 | 5.9 | 0.64 |
|  | KL08 | 78 ± 1.4 | 62 ± 3.8 | 85 | 62 | 82.2 ± 2.6 | 5.1 | 0.66 |
|  | IN09 | 87 ± 2.2 | 59 ± 3.2 | 92 | 66 | 86.7 ± 4.3 | 4.6 | 0.68 |

KL08=Directorate of Wheat Research (DWR) in 2008, Karnal, IN09=National Research Centre on Soybean
(NRCS) Indore in 2009, andDL10=Division of Genetics, Indian Agriculture Research Institute, New Delhi, India in

**Table 2.** Correlation analysis of quality related traits in RIL population derived from WL711/C306.

| Traits | GPC | SDS | HW | TKW | SD | WGC | DGC | FWA | DDT | DST | MTI | BDT | KH |
| --- | --- | --- | --- | --- | --- | --- | --- | --- | --- | --- | --- | --- | --- |
| **GPC** | 1 |  |  |  |  |  |  |  |  |  |  |  |  |
| **SDS** | 0.478* | 1 |  |  |  |  |  |  |  |  |  |  |  |
| **HW** | -0.354* | -0.214* | 1 |  |  |  |  |  |  |  |  |  |  |
| **TKW** | -0.687** | -0.178* | -0.485* | 1 |  |  |  |  |  |  |  |  |  |
| **SD** | -0.342* | -0.173* | -0.043 | -0.128* | 1 |  |  |  |  |  |  |  |  |
| **WGC** | 0.795** | 0.318** | -0.167* | 0.126* | 0.043 | 1 |  |  |  |  |  |  |  |
| **DGC** | 0.832** | 0.353** | -0.268* | 0.121* | 0.165* | 0.825** | 1 |  |  |  |  |  |  |
| **FWA** | 0.432** | -0.234* | -0.143* | 0.034 | -0.321* | -0.143* | 0.032 | 1 |  |  |  |  |  |
| **DDT** | -0.085 | 0.312** | 0.031 | 0.021 | 0.012 | 0.856** | -0.162 | 0.003 | 1 |  |  |  |  |
| **DST** | -0.041 | 0.483** | 0.023 | 0.124* | 0.114* | -0.243** | -0.173 | 0.008 | 0.881** | 1 |  |  |  |
| **MTI** | -0.108 | -0.323* | -0.284* | -0.542* | -0.013 | -0.092 | -0.101* | -0.343* | -0.583** | -0.777** | 1 |  |  |
| **BDT** | 0.095 | 0.014 | 0.053 | 0.154* | 0.003 | -0.226** | -0.155 | 0.021 | 0.822** | 0.877** | -0.767** | 1 |  |
| **KH** | -0.234* | -0.143* | 0.043 | 0.143* | 0.04 | 0.184* | 0.029 | 0.02 | 0.824** | 0.329* | 0.036 | 0.019 | 1 |

Grain protein content (GPC, %), Sedimentation rate (SDS), Hectolitre weight (HW, g), 1000-kernel weight (TKW, g), Moisture content (MC), Seed diameter (SD), Wet gluten content (WGC, %), Dry gluten content (DGC, %), Flour water absorption (FWA, %), Dough development time (DDT, min), Dough stability time (DST, min), Mixing tolerance index (MTI, F.U), Break down time (min) (BDT), Kernel hardness (KH)

### QTLs for bread making traits

38 putative QTLs related to 13 bread making quality traits were reported explaining 7.9% - 16.8% phenotypic variance (PV) by mapping analysis (Table 3, Fig 1). The QTLs were dispersed on 14 chromosome of all three (A, B and D) genome type i.e. 1A, 2A, 4A, 6A, 7A,1B, 3B, 4B, 7B, 1D, 2D, 3D, 4D and 5D. Six QTLs were identified for GPC on chromosome 1B, 1D, 3B, 3D, 5D and 7Aexplaining9.8% - 5.8% of PV. Alleles were contributed by WL71l at two QTLs (*qGPC.3D.1* and *qGPC.7A.1*) and from C306 at four QTLs (Table 3). The strongest effect for GPC (11.9) with 15.8 % PV was located on *qGPC.5D.1* with allele being contributed by C306. *qGPC.5D.1* was found to be co-located with the QTLs explaining HW, WGC, DST and KH. Another major QTL for GPC was *qGPC.7A.1* which explained 13.9% of PV and allele contributed by WL711. This QTL was observed to show co-location with QTLs responsible for SDS, TKW, DGC and KH.

**Figure 1:**
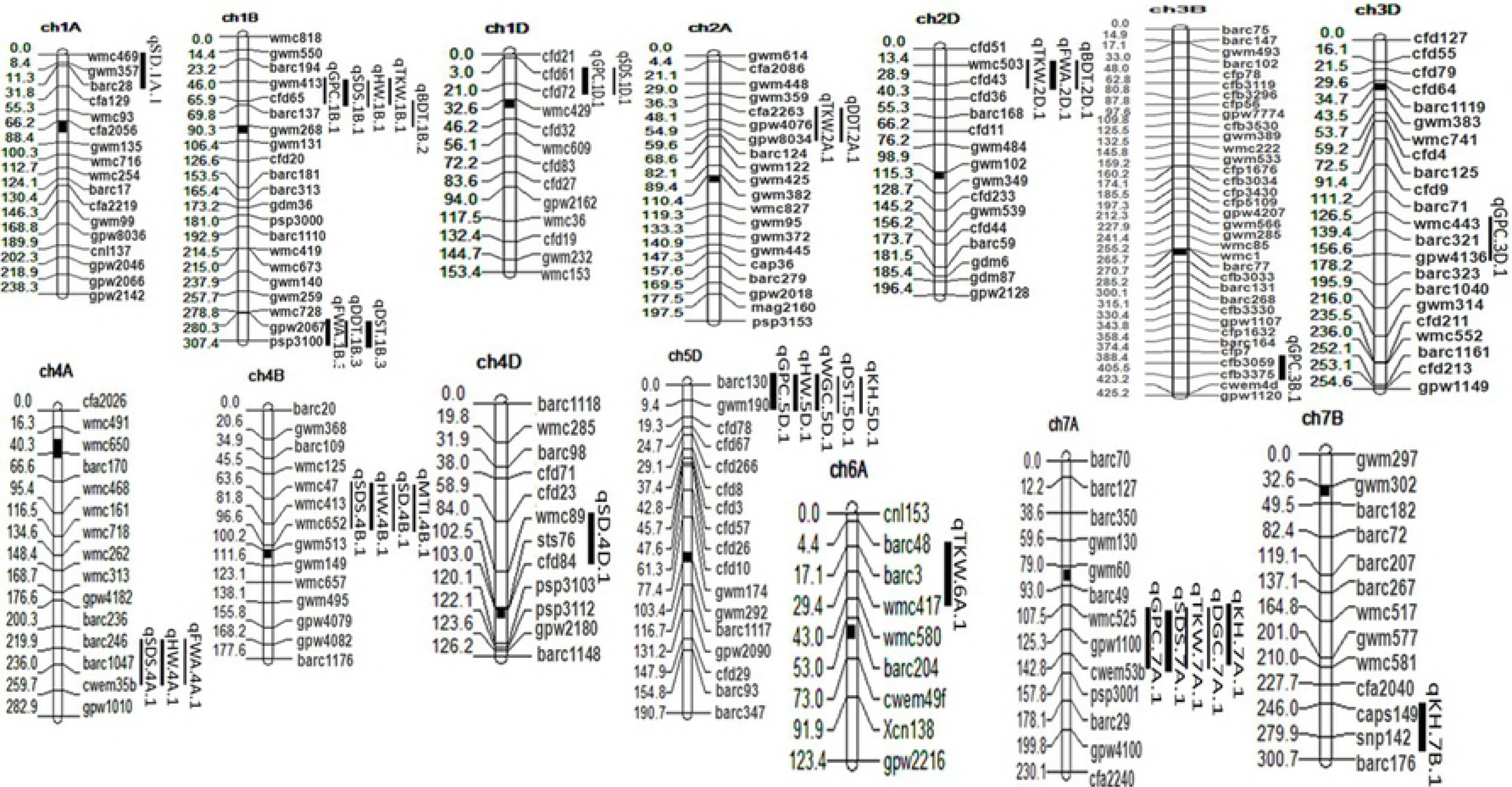
Quantitative trait loci (QTLs) for quality traits in WL711/C306 wheat RIL population. The vertical bars indicate the QTL confidence intervals. Map distances (cM) are shown on the left side of each chromosome.

**Table 3.** Major and minor QTLs for quality traits identified by composite interval mapping analysis using QTL cartographer software in 206 RIL population derived from WL711/C306.

| Traits<sup>1</sup> | QTL | Marker Interval | Position cM | LOD value | *a* Effect<sup>2</sup> | R<sup>2</sup>(%)<sup>3</sup> |
| --- | --- | --- | --- | --- | --- | --- |
| GPC | *qGPC.1B.1* | gwm413 - cfd65 | 43.7 | 3.1 | -4.7 | 10.3 |
|  | *q GPC.1D.1* | cfd61 - cfd72 | 2.6 | 4.3 | -3.6 | 12.4 |
|  | *qGPC.3B.1* | cfb3059 - cfb3375 | 317.1 | 5.7 | -2.6 | 12.8 |
|  | *qGPC.3D.1* | wmc443 - gpw4136 | 139.4 | 3.6 | 4.8 | 9.8 |
|  | *qGPC.5D.1* | barc130 - gwm190 | 0.01 | 7.8 | -11.9 | 15.8 |
|  | *qGPC.7A.1* | wmc525 - cwem53b | 115.5 | 3.1 | 6.9 | 13.9 |
| SDS | *qSDS.1B.1* | gwm413 - cfd65 | 43.7 | 4.5 | 34.6 | 11.6 |
|  | *qSDS.1D.1* | cfd61 - cfd72 | 2.5 | 3.3 | -2.3 | 10.1 |
|  | *qSDS.4A.1* | barc246 - cwem35b | 238.1 | 4.1 | 12.6 | 9.6 |
|  | *qSDS.4B.1* | wmc47 - wmc652 | 63.6 | 3.1 | 9.6 | 9.0 |
|  | *qSDS.7A.1* | wmc525 - cwem53b | 115.5 | 5.7 | -44.7 | 16.8 |
| HW | *qHW.5D.1* | barc130 - gwm190 | 0.01 | 7.4 | -46.1 | 14.7 |
|  | *qHW.4A.1* | barc246 - cwem35b | 238.1 | 3.8 | 5.7 | 10.5 |
|  | *qHW.1B.1* | gwm413 - cfd65 | 43.7 | 3.1 | 13.6 | 9.5 |
|  | *qHW.4B.1* | wmc47 - wmc652 | 63.6 | 3.6 | 4.7 | 11.6 |
| TKW | *qTKW.1B.1* | gwm413 - cfd65 | 43.7 | 3.5 | 26.2 | 10.6 |
|  | *qTKW.2A.1* | cfa2263 - gpw8034 | 54.9 | 3.6 | 28.6 | 10.3 |
|  | *qTKW.2D.1* | wmc503 - cfd43 | 28.9 | 4.3 | -37.9 | 10.8 |
|  | *qTKW.6A.1* | barc48 - wmc417 | 23.5 | 4.2 | -41.6 | 14.1 |
|  | *qTKW.7A.1* | wmc525 - cwem53b | 115.5 | 6.7 | -42.6 | 15.8 |
| SD | *qSD.1A.1* | wmc469 - barc28 | 8.4 | 3.7 | -2.1 | 12.6 |
|  | *qSD.4B.1* | wmc47 - wmc652 | 63.6 | 2.9 | 1.2 | 7.9 |
|  | *qSD.4D.1* | wmc89 - cfd84 | 102.5 | 5.8 | -2.6 | 14.7 |
| WGC | *qWGC.5D.1* | barc130 - gwm190 | 0.01 | 4.6 | -26.1 | 13.5 |
| DGC | *qDGC.7A.1* | wmc525 - cwem53b | 115.5 | 3.7 | -10.6 | 11.8 |
| FWA | *qFWA.1B.3* | gpw2067 - psp3100 | 292.3 | 4.2 | 63.2 | 14.1 |
|  | *qFWA.2D.1* | wmc503 - cfd43 | 28.9 | 4.3 | -36.8 | 15.3 |
|  | *qFWA.4A.1* | barc246 - cwem35b | 238.1 | 3.1 | 16.8 | 8.7 |
| DDT | *qDDT.1B.3* | gpw2067 - psp3100 | 292.3 | 4.1 | 2.6 | 11.4 |
|  | *qDDT.2A.1* | cfa2263 - gpw8034 | 54.9 | 3.6 | 1.8 | 10.3 |
| DST | *qDST.1B.3* | gpw2067 - psp3100 | 292.3 | 4.6 | -5.7 | 14.1 |
|  | *qDST.5D.1* | barc130 - gwm190 | 0.01 | 3.2 | -3.8 | 10.4 |
| MTI | *qMTI.4B.1* | wmc47 - wmc652 | 63.6 | 3.1 | 54.6 | 8.7 |
| BDT | *qBDT.1B.2* | cfd65 - gwm268 | 77.8 | 3.5 | 12.5 | 9.4 |
|  | *qBDT.2D.1* | wmc503 - cfd43 | 28.9 | 4.3 | 11.5 | 10.8 |
| KH | *qKH.5D.1* | barc130 - gwm190 | 0.01 | 3.8 | 46.8 | 9.5 |
|  | *qKH.7A.1* | wmc525 - cwem53b | 115.5 | 5.4 | -65.4 | 14.6 |
|  | *qKH.7B.1* | caps149 - snp142 | 256 | 4.1 | -55.4 | 13.6 |

Grain protein content (GPC, %), Sedimentation rate (SDS), Hectolitre weight (HW, g), 1000-kernel weight (TGW, g), Seed diameter (SD), Wet gluten content (WGC, %), Dry gluten content (DGC, %), Flour water absorption (FWA, %), Dough development time (DDT, min), Dough
stability time (DST, min), Mixing tolerance index (MTI, F.U), Break down time (BDT, min), Kernel hardness (KH)
2, 3 additive main effects, R<sup>2</sup> (*a*) % phenotypic variation explained by *a* effects. A positive value of the additive main effects (*a*) indicates that WL711 contributes allele to increase the trait, and a negative value means that C306 provides allele to increase the trait.

Five QTLs associated with SDS sedimentation were identified on chromosome 1B, 1D, 4A, 4B and 7A explaining 9.0 - 16.8% of PV. Out of the five QTLs for SDS sedimentation, three QTLs (*qSDS.1B.1*, *qSDS.4A.1* and *qSDS.4B.1*) were coming from WL711 and two (*qSDS.1D1* and *qSDS.7A.1*) were from C306 (Table 3). The largest favourable effect on SDS sedimentation (44.7) was associated with *qSDS.7A.1* explaining 16.8% of PV, allele contributed by C306. Another major impact QTL for SDS qSDS.1B.1with major additive effect (34.6) was identified on chromosome 1B explaining 11.6% of PV.

Four QTLs for HW were reported on chromosomes 1B, 4A, 4B and 5D explaining 9.5% - 14.7% of PV with three QTLs were coming from WL711 and one from C306 (Table 3).A QTL *qHW.5D.1* having additive effect (46.1) and explaining 14.7% of PV and co-localized with QTLs for GPC, WGC, DST and HW. For TKW total 5 QTLs were reported located on chromosomes 1B, 2A, 2D, 6A and 7A explaining 10.3% - 15.8% of PV. A major QTL, *qTKW.7A.1* for TKW was identified on 7A with allele coming from C306 explaining 15.8% of PV. This QTL was co-located with QTL for GPC, SDS, DGC and KH.

Three QTLs were identified for SD on chromosomes 1A, 4B and 4D explaining 7.9% - 14.7% of PV. Another QTL for SD was identified on 4D, *qSD.4D.1* explaining 14.7% of PV, positive allele was contributed by C306 (Table 3). Another major QTL responsible for WGC was identified on chromosome 5D explaining 13.5% of PV in which positive allele was contributed by C306. A major QTL *qDGC.7A.1*, for DGC was identified on chromosome 7A explaining 11.8% of PV, positive allele was contributed by C306 and was found to be co-located with QTLs for GPC, TKW and KH. Total of 3 QTLs for FWA were identified on chromosomes 1B, 2D and 4A explaining 8.7% - 15.3% of PV. A major QTL, *qFWA.2D.1* was identified on 2D explaining 15.3% of PV with positive allele contribution by C306. This QTL co-located with QTL for TKW and BDT. Two QTLs for DDT were identified on chromosomes 1B and 2A. Another QTL for DDT was identified on 2A, *qDDT.2A.1* explaining 10.3% of PV and positive allele contribution done by WL711 and co-located with TKW. Two QTLs namely *qDST.1B.3* and *qDST.5D.1* identified for DST were located on chromosomes 1B and 5D explaining 14.1 and 10.4% of PV respectively. A major QTL for DST, *qDST.1B.3* on chromosome 1B explaining 14.1% of PV with allele contribution by C306.

A minor QTL responsible for MTI was identified on 4B explaining 8.7% of PV. Two QTLs for BDT were identified on chromosomes 1B (*qBDT.1B.2*) and 2D (*qBDT.2D.1*) explaining 9.4 and 10.8% of PV respectively (Table 3). Two major QTLs and a minor QTL for KH was identified on chromosomes on 7A, 7B and 5D explaining 9.5 to 14.6% of PV. A major QTL, *qKH.7A.1* for KH on chromosome 7A explaining 14.6% of PV and positive allele contributed by C306.

### QTL x environment interactions and Epistatic QTL

The effects of the QTL x environment interactions (QE) for quality related traits were recorded and listed in Table 4. From the measured quality traits, GPC and TKW two QQ interactions were detected. In addition, few genomic regions identified in this study showed QE, QQ and QQE interactions and their effects were very less noticeable than the main additive effects (a). Indicated that the additive effects were more significant in comparison with epistatic effects in the studied quality traits. Epistatic QTLs showed QTL x QTL (QQ) and QTL x QTL x environment (QQE) interaction.

**Table 4.** Epistatic QTLs and QTL x QTL x environment interaction for quality related traits identified by two locus analysis using QTL Network software in 206 RILs derived from WL711/C306.

| Traits<sup>a</sup> | QTL_i<sup>b</sup> | Interval_i | QTL_j | Interval_j | QQ<sup>c</sup> | QQE<sup>d</sup> | R<sup>2</sup> % |  |
| --- | --- | --- | --- | --- | --- | --- | --- | --- |
|  |  |  |  |  |  |  | QQ<sup>e</sup> | QQ E<sup>f</sup> |
| GPC | *qGPC.1B.1* | gwm413 - cfd65 | *qGPC.1D.1* | cfd61 - cfd72 | 0.21 | 0.16 | 0.08 | 0.03 |
| TKW | *qTKW.1B.1* | gwm413 - cfd65 | *qTKW.2D.1* | wmc503 - cfd43 | -0.42 | -0.13 | 0.08 | 0.05 |

A positive value means that the parent-type effect is greater than the recombinant-type effect
A negative value means that the parent-type effect is less than the recombinant-type effect
a GPC- grain protein content, TKW- thousand kernel weight
bQTL\_i and QTL\_j are a pair of QTL involved in epistasis
c QQ, the epistatic main effect d QQE, the epistasis x environment interaction effects
e R<sup>2</sup> (QQ) %, Phenotypic variation explained by QQ effects
f R<sup>2</sup> (QQE) %, Phenotypic variation explained by QQE effects

### Candidate genes identification by using syntenic analysis

Identified QTL regions for GPC located on 1BS chromosome of wheat was compared with rice genomic regions. BLAST function was performed using DNA sequences of respective marker sequences. Total of 33.3% sequence homology detected (e-value ≤10^-20^) with the rice genomic sequences and rice candidate gene namely *GLU1* for glutenin was identified on wheat chromosome 1BS (Table 5).

**Table 5.** Orthologous relationship between wheat DNA sequences and rice genes on chromosomes 1BS and 5DS. The table displays the result of BLASTN analysis against the rice allocated genes (TIGR Rice Annotation Release 5;http://rice.tigr.org/). For each candidate gene, the percentage of identity, length of the alignment, co-ordinates of the wheat (query) and rice (subject) best hit sequences, E value and score are given.

| QTLs | Rice Gene Id | Putative function | % Identity | Match length | Query Start | Query End | Subject Start | Subject End | E-value | Score |
| --- | --- | --- | --- | --- | --- | --- | --- | --- | --- | --- |
| *qGPC.1D.1* | <u>Os02g0249600</u> | GLU1 (GLUT ELIN 1) | 73.16 | 59 | 349 | 475 | 389 | 493 | 1.00E-25 | 289 |
| *qKH.5D.1* | <u>Os02g0743400</u> | PIN1A (PIN protein 1A) | 78.42 | 47 | 284 | 491 | 435 | 524 | 1.00E-43 | 415 |

On wheat chromosome 5DS, BLAST function was performed using DNA sequences of respective marker sequences. Total of 33.3% sequence homology detected (e-value ≤10^-20^) with blast region of the rice genomic sequences and rice orthologous gene namely *PIN1A* was probable candidate gene found on wheat chromosome 5DS (Table 5).

## Discussion

Growing genotypes under well adapted conditions with strong phenotypic expression can lead to over estimationof the genetic component and it could be avoided by including contrasting environments and seasons in which observations are made. In accordance the experimental materials consisting of 206RIL population developed from the cross WL711/ C306 were grown under three environmental condition. Total of 38 QTLs have been identified through CIM for thirteenquality related traits across environments. Continuous phenotypic variation and transgressive segregation for all the traits have been observed in the RIL population revealed the quantitative inheritance of these traits. Further, presence of alleles from both the parentsand usefulness of this populationfor QTL analysis.

Increased GPC is focus area of current wheat quality breeding program. It is broadly recognised that GPC is highly reliant on environment, with high protein when stress condition occurred during the grain filling stage. Similarly, parent C306 and the RILs showed mean GPC significantly high (above 15%) in the IN09 environment, where RILs were exposed to heat during post anthesis to grain filling stage. These results were in agreement with Maphosa *et al.(25*). GPC showed low value (below 12%) in the DL10 and KL08 experiment, when crop experienced cool and moist conditions. Li *et al*. (26) indicated that total GPC is linked to temperature and low humidity. The negative correlation between GPC and TKW was recorded in this population which was distinguished in previous studies as well (27). The QTLs related to GPC were reported earlier on the regions of several chromosomes showing several loci controlling wheat GPC, those studies also suggested very less differences in GPC in parental line, had QTLs were still detected(28, 29). In present study, QTL analysis for GPC revealed six QTLs with PV ranging from 9.8% -15.8% located on six different chromosomes i.e. 1B, 1D, 3B, 3D, 5D and 7A. Though the difference in protein content between the parents was less, transgressive segregants were observed for GPC. These transgressive segregants for high GPC might be due to minor genes segregating in the population and the different GPC controlling alleles in the parents, confirming the suitability of this population for QTL analysis for GPC (Chee et al.2001). A set of epistatic QTL showed weak additive × additive × environment effects (AAE) interaction suggested additive effects played a most important role in wheat GPC.

In this study, QTLs for GPC and SDS were mapped near to *Glu*-*D1*region present on the chromosome 1D which is similar to the results observed in other studies as well (31, 32). In fact, *Glu*-*D1*gene coded HMW subunits (2+12 and5+10) were found to affect the protein quality in ChSh population (33). Furthermore another wheat protein triticin at *Tri*–*D1* was reported to be positively affecting wheat dough bread making quality which was also present on the short arm of chromosome 1D (34). When compared with the genomic DNA sequences of rice syntenic to wheat chromosome 1DS candidate gene namely *Glu1* for glutelin was identified(Table 5). The *Glu1* locus identifies as a clusters of storage polypeptides named high molecular weight (HMW) subunits of glutenin (35). The other two QTLs for GPC on chromosomes 3B and 5D had larger effects, also present chances for useful genetic improvement.

TKW is one of the important yield components and selection for this trait directly increases the yield (36). Though its correlation with quality parameters is reported (37). Selection for quality trait alone will not help in improving this trait. A pronounced and significant variation for TKW suggested several genes with major and minor effects that were involved in the phenotypic expression of this trait. TKW was controlled by 5 QTLs identified in our study which were present on the chromosomes -1B, 2A, 2D, 6A and 7A. Sun *et al*. (38) also identified seven QTL regions and contributing chromosomes were reported to be 2A, 3B, 4A, 5D, 6A, 6B, and 7B in RIL population. In addition, Reif *et al*. (39) identified 12 putative QTLs on genomic regions of chromosomes 1A, 3A, 5A, 7A, 1B, 3B, 6B, 1D, 3D, 4D, and 7D in RIL population using association mapping. In these two studies only single QTL on chromosome 7A was similar which suggested that many genes governing the trait TW. Sun *et al*. (38) identified eight QTL in Ning7840 × Clark RIL population present in chromosomes 1D, 2D, 4A, 4B, 5A, 5A, 5B and 6A but only 2D and 6A were present in this study. Similarly, three novel QTLs were detected on chromosomal regions of 1B, 2A, and 7A in the present study. In this study, one epistatic QTL was identified with non-significant Additive × environment (AE) or AAE interactions which showed that an additive effect responsible for the main genetic variance of TKW.

KH played a major role for determining quality of bread wheat and end use properties. Also, *Ha* locus mainly known for affecting grain hardness in wheat. Several QTLs for KH have been reported in different mapping population that distributed on all over twenty one wheat chromosomes except for 3D and 6A (40). Earlier studies have also identified QTLs for KH on each of these chromosomes (41). Both parents contributed favourable alleles for KH which confirmed the quantitative nature of the trait. Comparative analysis with rice genomic DNA sequences with wheat chromosome 5DS candidate gene namely *PIN1A* was identified (Table 5). In wheat, associations of qualitatively inherited genes together represent gene-rich regions and they form the hot spots of recombination. QTL are usually spread over all the chromosomes, but clusters of QTL in certain chromosomal regions have been observed.QTL affecting several traits are common and may be due to pleiotropy or close linkage (32). Since most of the QTL hotspots in this study were located in the short and long armof the chromosomes, QTL co-location of yield QTLs have alsobeen identified previously in wheat (1,36). Similarly, 5 QTLs were mapped on 5D and 5 QTLs on 7B chromosome and 4 QTLs on 1B in which some of them showed stability across the environments also suggested that two QTL clusters supposed to have pleiotropic effects. It is likely that they represented similar genes/protein content. Several linked markers in the clusters suggesting the usefulness of these markers for marker-assisted breeding of these QTLs to enhanceend product quality of wheat.

## Conclusions

38 QTLs for total 13 end product quality traits were studied explaining 7.9 (*qSDS.4B.1*) to 16.8 (*qSDS.7A.1*) of PV detected on 14 chromosomes i.e. 1(ABD), 2(A, D), 3(B,D), 4(ABD), 5D, 6A, 7A and 7B. The additive effect was found to be positive in 17 QTLs and was contributed by WL711 and 21 were negative and contributed by C306. Total eight QTLs for three major quality traits affecting bread making quality namely SDS (5), DST (2) and DGC (1) were identified with 9.6 to 16.8 % PV. For SDS, from five of the three alleles were contributed by WL711, for DST both were contributed by C306 and DGC alleles were contributed by C306. For GPC, six QTLs were reported on chromosome 1B, 1D, 3B, 3D, 5D and 7A showing 9.8%-15.8% of PV for the trait with positive allele coming from WL71l at two QTLs (*qGPC.3D.1* and *qGPC.7A.1*) and from C306 at four QTLs. The strongest effect for GPC (11.9) with 15.8 % PV was located on *qGPC.5D.1* with positive allele being contributed by C306. This study revealed the importance of combination of stable QTL with region specific QTL for better phenotype and QTLs presented in our study will be useful in MAS efforts for improvement of wheat grain and bread making quality.

## Acknowledgements

The authors would like to thank the Indian council of Agricultural Research (ICAR) for supporting this work. The authors would also like to extend their sincere appreciation to the Deanship of Scientific Research at King Saud University for its funding to the Research Group number (RG-1435-014).

